# Developmental regulation of oocyte lipid uptake via ‘patent’ follicular epithelium in *Drosophila melanogaster*

**DOI:** 10.1101/2019.12.31.891838

**Authors:** Sarayu Row, Wu-Min Deng

**Affiliations:** Department of Biological Science, Florida State University, Tallahassee, FL 32306-4295, USA; Department of Biochemistry and Molecular Biology, Tulane University School of Medicine, New Orleans, LA 70112, USA

## Abstract

Epithelia form protective permeability barriers that selectively allow the exchange of material while maintaining tissue integrity under extreme mechanical, chemical, and bacterial loads. Here, we report in the *Drosophila* follicular epithelium a developmentally regulated and evolutionarily conserved process, ‘patency’, wherein a breach is created in the epithelium at tricellular contacts during mid-vitellogenesis. In *Drosophila*, patency exhibits a strict temporal range delimited by the transcription factor Tramtrack69, and a spatial pattern regulated by the dorsal-anterior signals of the follicular epithelium. Crucial for lipid uptake by the oocyte, patency is also exploited by endosymbionts such as *Spiroplasma pulsonii*. Our findings reveal an evolutionarily conserved non-typical epithelial function in a classic model system.

## Introduction

Epithelial integrity is maintained through cell division, apoptosis, and morphogenetic movements *(1,2)*, and the preservation of tricellular contacts (TCs) is key to its physical and electrochemical barrier function *(3)*. Failure to maintain TCs can disrupt epithelial integrity and have catastrophic consequences such as metastatic cancer *(4,5)* and Crohn’s disease *(6)*. The epithelium is also permeable, allowing selective passage of material via transcellular and paracellular routes *(7)*. The extensively characterized *Drosophila* follicular epithelium (FE) surrounding developing egg chambers provides an excellent model to study the balance between barrier function and permeability in epithelia. As the developmental unit of oogenesis, each egg chamber is comprised of an oocyte and 15 nurse cells, all enclosed in the FE monolayer *(8)*. A string of sequentially developing egg chambers beginning with the germarium and advancing from stages 1 through 14 is contained in each of the 16 ovarioles in each ‘meroistic’ ovary *(8)* [Fig. 8.]. The FE monolayer protects and supports germline development while going through a series of spatiotemporally regulated morphogenetic movements, and also secretes the chorion, or eggshell *(8, 9)*. Vitellogenic stages (stage (St) 8-12) are characterized by the trans-epithelial movement of hemolymph-borne yolk proteins (YPs) into the oocyte. YPs are synthesized primarily in the fat body, secreted into the hemolymph, and travel between the follicle cells to reach the oocyte membrane, and are internalized via receptor-mediated endocytosis *(10–12)*. The FE also synthesizes YPs in small amounts until St 11 when it switches to chorion secretion *(13, 14)*. Other materials carried by the hemolymph have been postulated to enter the oocyte, such as lipophorins and endosymbionts *(15,16)*, but the mechanistic details of trans-epithelial FE transport remain unclear.

In this study, we present a mechanism for the trans-epithelial transport of lipids across the FE, a phenomenon termed ‘patency’, previously unreported in *Drosophila*. Unlike in other insects, we found that patency in *Drosophila* is only present in mid-vitellogenesis. This reduced temporal range appears to be regulated by the BTB transcription factor Tramtrack69 (Ttk69). We also report a consistent spatial pattern of patency in the FE, influenced by the signaling pathways that pattern the dorsal anterior. Moreover, we found the spatiotemporal patterns of patency are evolutionarily conserved in the *Drosophila* species. Together, our results reveal a developmentally regulated function of the follicular epithelial tissues, wherein a breach is created for trans-epithelial material transfer.

## Results

### *Drosophila* follicular epithelium exhibits patency

While examining wildtype (WT) *Drosophila melanogaster* egg chambers, we noticed an anomalous but consistent feature — the opening of the TCs in mid-vitellogenic FE. These TC ‘gaps’ are present at the basal, lateral, and apical domains of the FE [Fig. 1(A-D”), Fig. S1(B), (D-H)], effectively creating a breach in the epithelium. Although not previously described in *Drosophila*, several insect species have been reported to exhibit ‘patency’, a phenomenon characterized by the opening of intercellular contacts between follicle cells for oocyte yolk uptake during vitellogenesis *(17)*. We confirmed the appearance of TC gaps using a milder detergent-free immunostaining protocol, as well as electron microscopy [Fig. S1(A-A”), Fig. 1(E-E’)], demonstrating that *Drosophila* FE indeed exhibits patency.

**Fig. 1.**
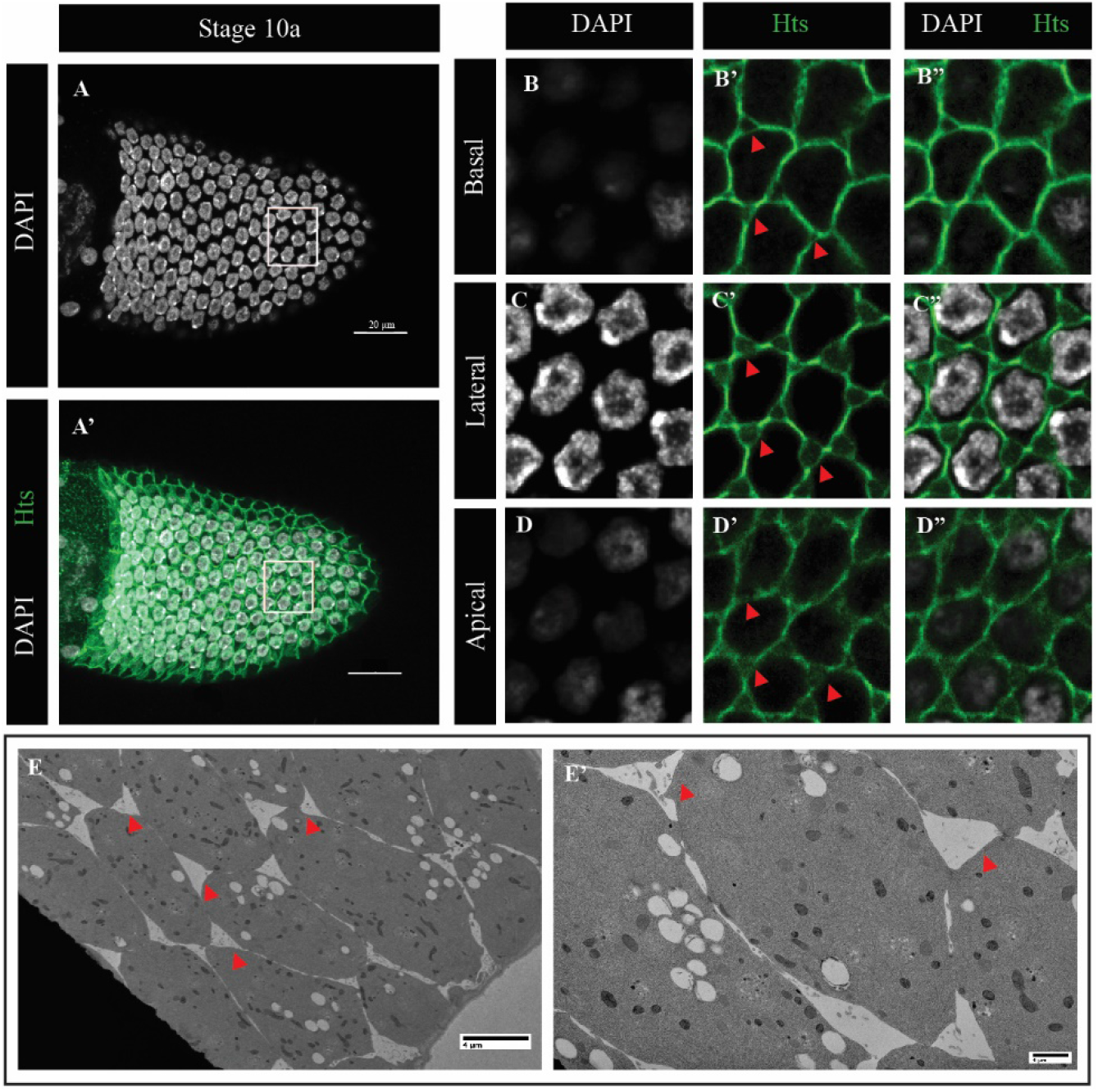
Patency in *Drosophila melanogaster.* (A-A’) Stage 10a oocyte-associated FE. White box is expanded in (B-D”). (B-B”, C-C”, D-D”) Basal, lateral, and apica l view, respectively, of the TC gaps in oocyte associated FE at St 10a. Hts (Hu Ii tai shao, adducin) marks the membrane, DAPI marks the nuclei. (E-E’) TEM images of the surface of a St10a FE. TC gaps indicated by red arrowheads.

### Temporal range of patency and regulation by Ttk69

Unlike in other insects (e.g., *R. proxilus*) where patency is present in most vitellogenic stages *(18)*, we found that patent TCs in *Drosophila* FE are present only during mid-vitellogenic stages (St10a and 10b), suggesting a more limited temporal range [Fig. 2(A-E”)]. Immunostaining with various epithelial markers revealed a dynamic pattern of junction and adhesion proteins in the TCs over the course of patency [Fig. S2(A-T”)]. Septate junctions, marked by Discs large (Dlg), lose their connections at TCs during patent stages and are reconnected at stage 11 when the gaps disappear [Fig. S2(A-D”)]. Adherens junctions, represented by E-cadherin (E-cad), are removed from TCs in patent epithelia, and reappear when patency is terminated [Fig. S2(E-H”)]. In contrast, cortical F-actin remains intact and continuous through mid-vitellogenesis, although marginally reduced, indicating that the structural integrity might not be compromised even in patent FE [Fig. S2(I-L”)]. The tricellular junction (TCJ) septate junction protein Gliotactin (Gli) *(19)*, however, only appears at the TCs starting at late stage 10b [Fig. S2(M-P”)], coincident with the cessation of patency. The accumulation of Gli at TCs suggests the assembly of the TCJs - at least the tricellular septate junctions - at the termination of patency, tipping the functional balance of the FE away from trans-epithelial transport, and in favor of barrier function for the remainder of oogenesis. Additionally, the extracellular matrix protein Laminin accumulates in the gaps at St 10b [Fig. S2(Q-T”’)], indicating the possibility of basement membrane components also being transported across the epithelium, perhaps to aid in the closing or maintenance of the TCs post-patency. Consistent staging of egg chambers was based on Jia *et al*., 2015 *(20)*.

**Fig. 2.**
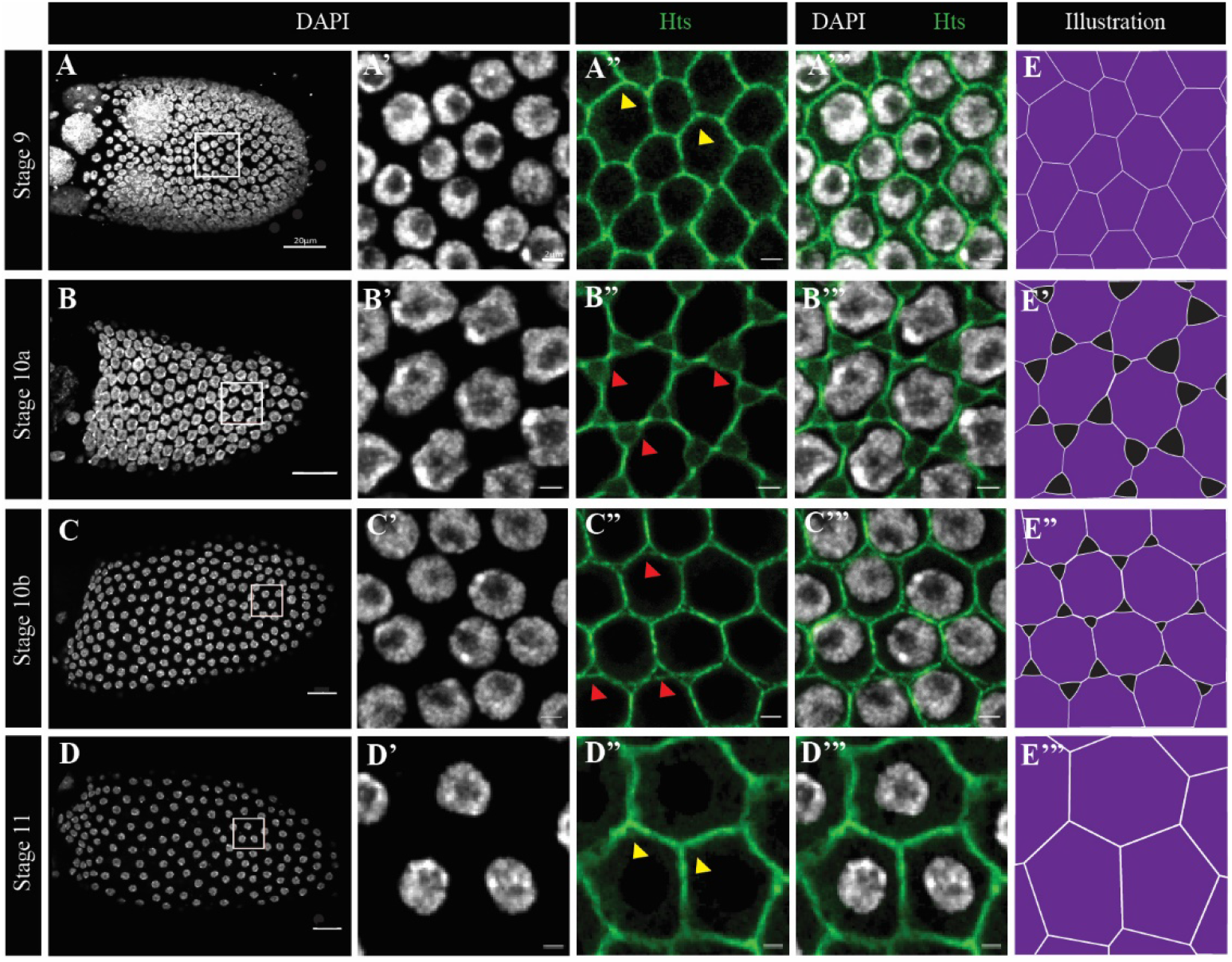
Temporal range of patency. (A-D) Projections ofSt9-Sll 1 egg chambers. Regions in the white box are expanded in (A’-A”’ to D’-D”’). Intact res at stages 9 and 11 (yellow arrowheads in A” and D” respec­ tively), and patent res at stages 10a and 1Ob (red arrowheads in B” and e” respectively).(E-E”’) Illustra­ tions of FE at stages 9 through 11 depicting intact and patent res. Hts marks the membrane (green) and DAPI marks nuclei.

Next, to determine how patency is temporally regulated, we performed an RNAi screen for transcription factors known to be active during mid-oogenesis, and identified Ttk69 - a zinc-finger transcription factor that coordinates FE morphogenesis by regulating the expression levels and localization of adhesion proteins such as E-Cad *(21, 22)*. Ttk69, previously reported to be expressed starting at St 10a, is also expressed during pre- and early vitellogenic stages [Fig. S3(A-D’)], and knocking down *ttk69* resulted in a larger temporal range of patency, with ectopic TC gaps ranging from St 9-11 [Fig. 3(A-D””)]. Similar to normal patent FE [Fig. S2(F-G”)], *ttk69*-knockdown FE with premature patency exhibited a lack of E-cad at TCs [Fig. S3(E-F’”)], suggesting that Ttk69 plays a role in restricting the temporal range of patency in the FE.

**Fig. 3.**
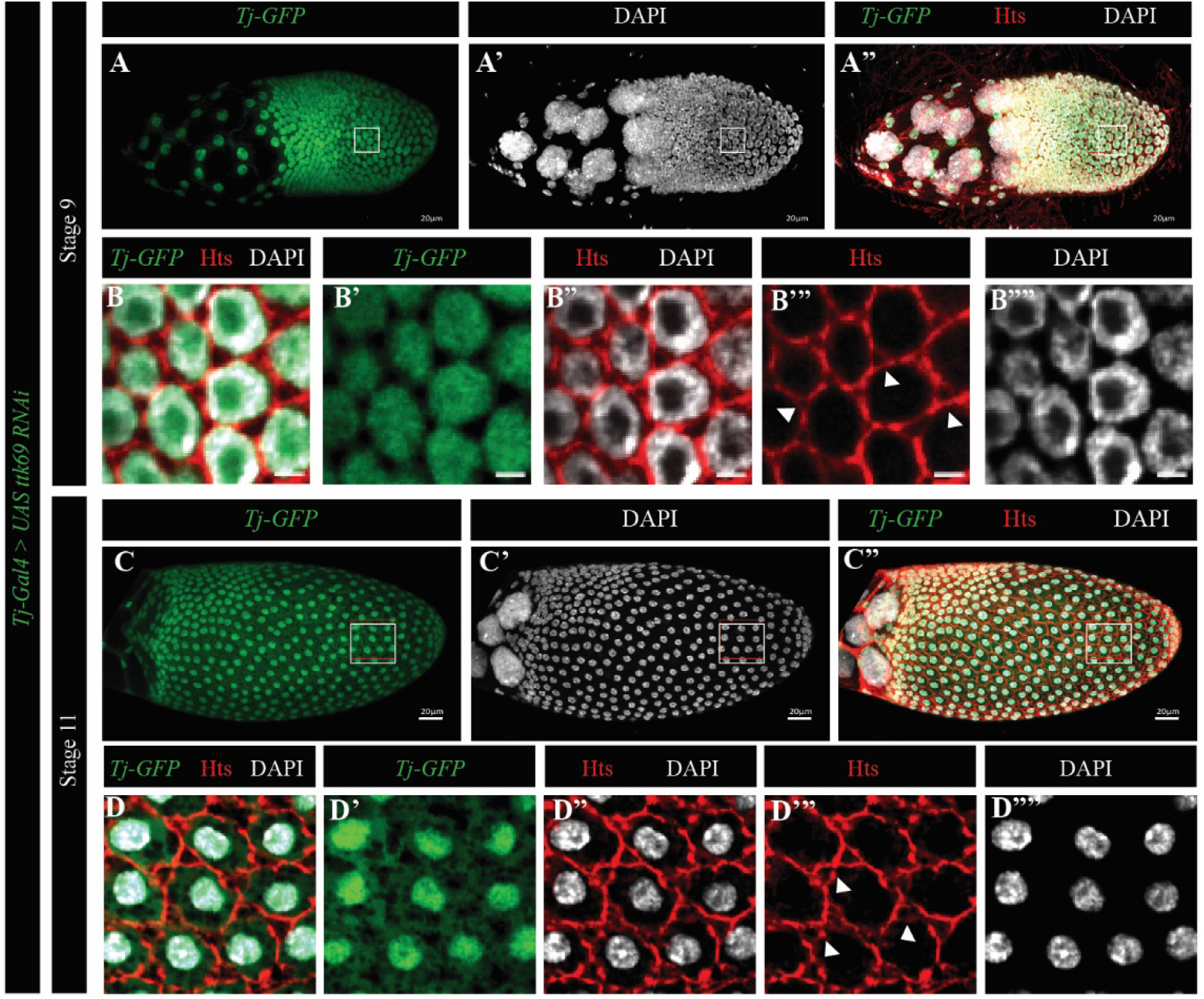
Ttk69 regulates the temporal range of patency. (A-D””) *UAS-ttk69-RNAi* expressing FE under *TJ> Gal4,* marked by *UAS-GFP.* ttk69 KD St 9 (A-A”) and St 11 (C-C”) egg chambers; white boxes are enlarged in (B-B””) and (D-D””) respectively, both showing ectopic gaps (white arrowheads). Hts marks the membrane (red), and DAPI marks nuclei.

### Spatial pattern of patency is influenced by the dorsal anterior signals of the FE

While examining St 10b egg chambers, we discovered that patency also shows a spatial pattern in the FE - specifically, the dorsal-anterior region has intact TCs while the rest of the FE TCs are patent at St 10b [Fig. 4(A-B’”, G-G’)]. The *Drosophila* FE is patterned along the anteroposterior and dorsoventral axes by the BMP (Bone Morphogenetic Protein) or Decapentaplegic (Dpp), and Epidermal Growth Factor Receptor (EGFR) pathways, creating a well-characterized subset of cells in the dorsal anterior with elevated levels of adhesion proteins (eg., E-cad, Fasciclin 3 (Fas3)) *(23–25)* [Fig. S4(A-B’”)]. Ectopic expression of Dpp in the whole FE resulted in elevated E-cad and Fas3 levels across the FE [Fig. S4(F-G”)], and all TCs remained intact at St 10a [Fig. 4(C-D”, H-H’)]. In contrast, removal of EGF signaling by expressing a dominant negative form of EGFR in the FE resulted in the loss of E-cad and Fas3, and ectopic patency in the dorsal anterior FE, thus eliminating the spatial pattern of patency [Fig. S4(H-I”), Fig. 4(E-F”, I-I’)]. Together, our data indicate that the spatial distribution of patency in the FE is largely regulated by Dpp and EGFR signaling.

**Fig. 4.**
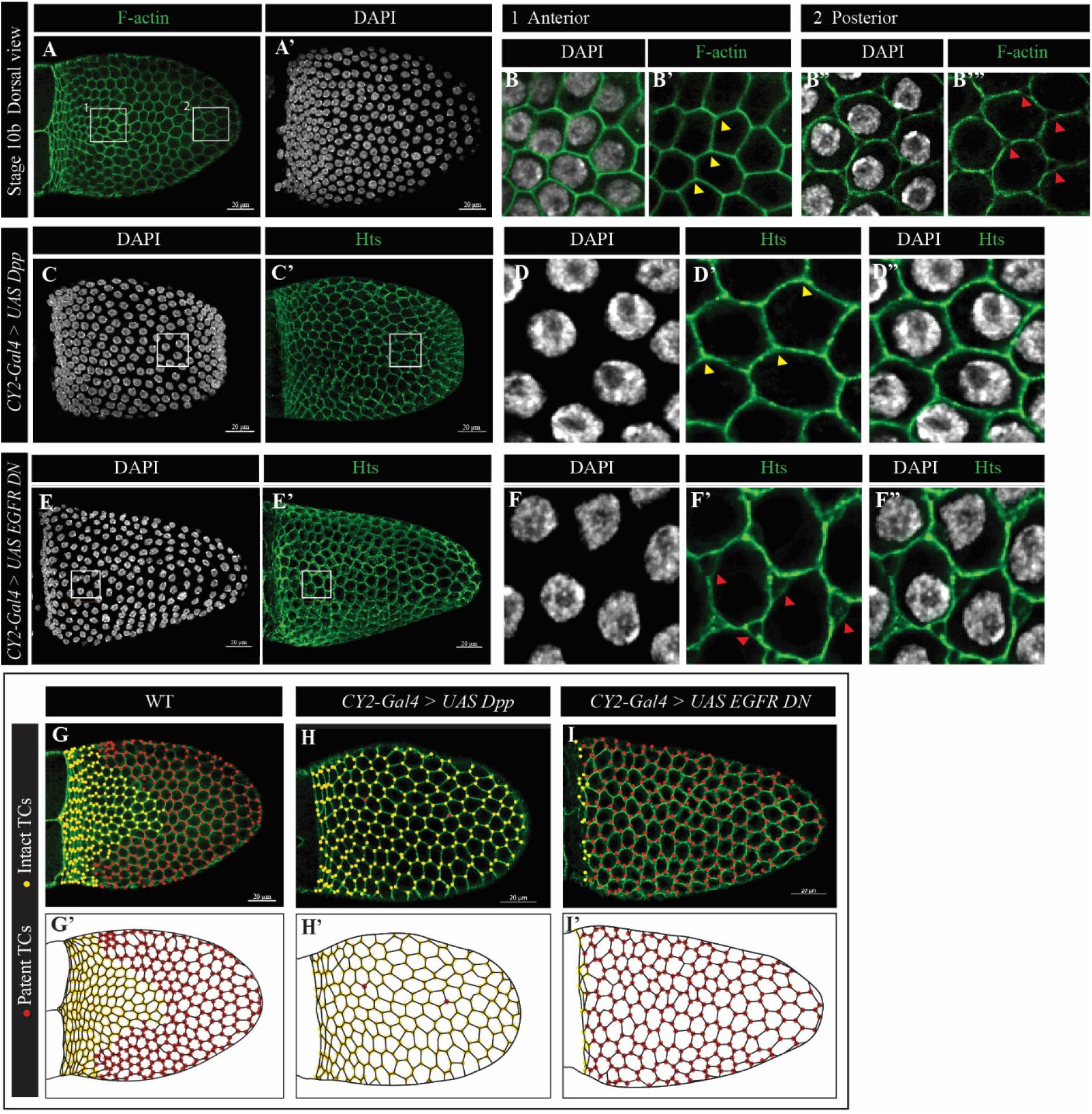
Spatial pattern of follicular patency and regulation. (A-A”) Dorsal view ofa St 1Ob egg chamber. Box#l in the dorsal-anterior is enlarged in (B-B’) showing intact TCs (yellow arrowheads); box#2 in the dorsal posterior is enlarged in (B”-B’”) showing patent TCs (red arrowheads). (C-C’) Dorsal view of a *UAS-Dpp* expressing FE. White box is enlarged in (D-D”) showing absence of patency (yellow arrowheads) (E-E’) Dorsal view of *UAS-EGFR-DN* expressing FE. White box is enlarged in (F-F”) showing ectopic patency in the dorsal anterior (directly above the oocyte nucleus) (red arrowheads). (G-1’) Illustration of the spatial pattern of patency, with yellow dots marking intact TCs, and red dots marking patent TCs in WT (G-G’), *UAS-Dpp*(H-H’), and *UAS-EGFR-DN (l-1’)* expressing FE.

### Patency is required for oocyte lipid uptake, and assists in vertical transfer

During oogenesis, lipid levels show a marked increase in the oocyte at St 10a *(26)*, which, incidentally, is also when TC gaps appear in the FE, and we therefore hypothesized that lipids are transferred across the FE into the oocyte via patency. Indeed, our TEM pictures showed material that could possibly be lipids in the gaps of patent FE [Fig. S1(C)]. Further investigation using neutral lipid dyes Nile red and BODIPY revealed that lipids were in fact present in the TC gaps, and appeared to be moving across the epithelium through the gaps [Fig. 5(A-C””)]. Additionally, in egg chambers with FE lacking patency, the oocyte lipid levels were significantly reduced at St 10a [Fig. 5(D-E”)], suggesting that follicular patency is necessary for oocyte lipid uptake. Consistently, egg chambers which develop with FE lacking patency appear to be abnormal, with a smaller oocyte at later stages, in addition to the expected lack of dorsal appendages as a consequence of ectopic *UAS-Dpp* expression [Fig. 5(F-F’)].

**Fig. 5.**
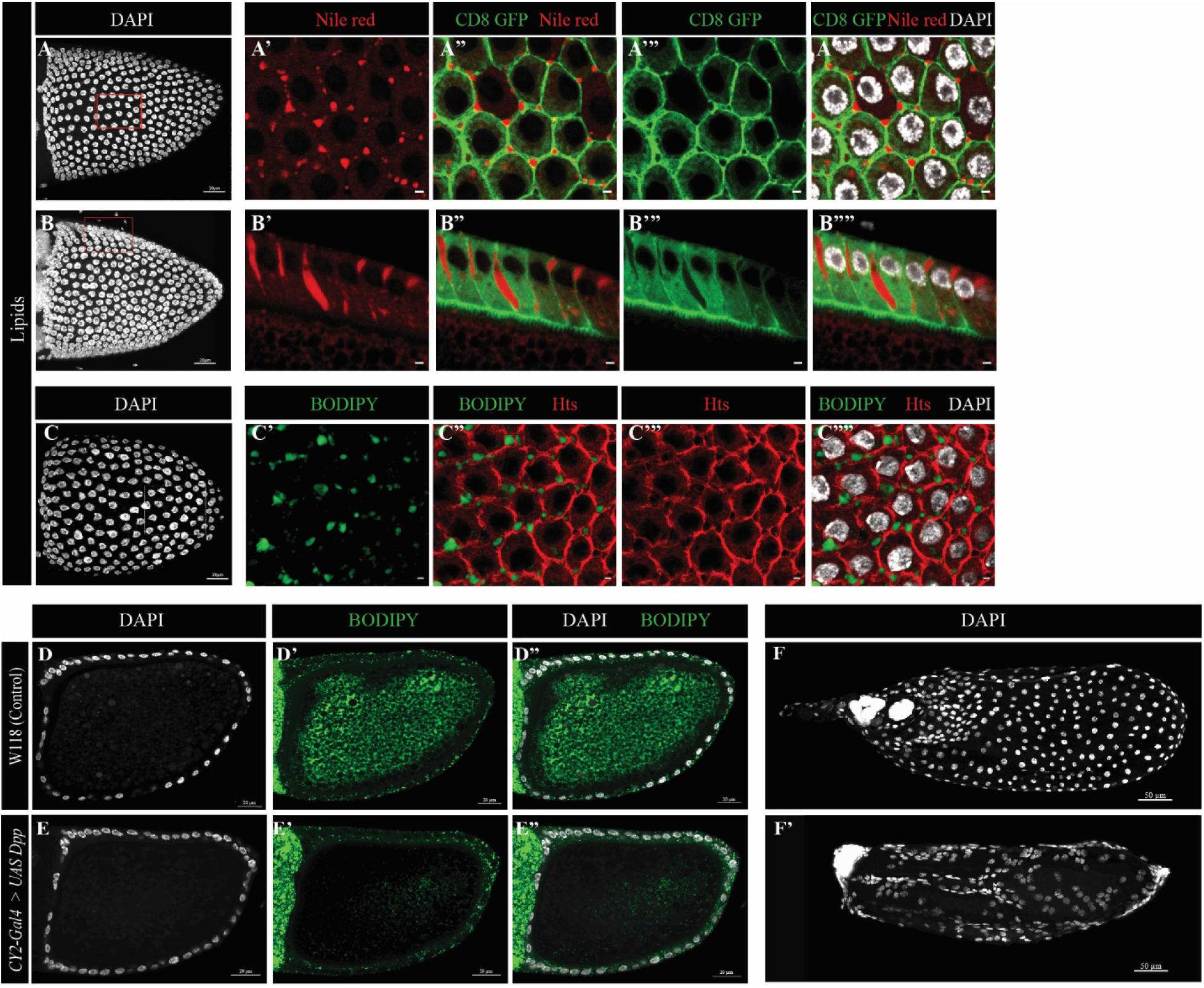
Patency for lipid uptake by the oocyte at St 10a. (A-C””) Lipids are present in the gaps in patent FE. (A-B””) Nile red staining shows lipids in the TC gaps, and spanning the FE. Basal and cross section views (A’-A””, B’-B”” respectively). CD8-GFP marks the membrane. (C-C””) BODIPY493/503 (green) confirms presence of lipids in the gaps. Membrane is marked by Hts (red). BODTPY sta ining of control WT St 1Oa egg chamber (D-D”) and *UAS-Dpp* expressing FE lacking patency showing reduced le ve ls of oocyte lipi ds at St IOa (E-E”). Note FE lipid glo bules are present in both. (F-F’) Later stage egg chambers with (F) WT FE and (F’) *UAS-Dpp* expressing FE, resulting in abnomial egg chambers. DAPT marks the nuclei.

Recently, the hormone ecdysone was shown to induce lipid uptake at St 10a by an unknown mechanism *(26)*. When we expressed a dominant negative form of the ecdysone receptor EcR B1, the FE showed no signs of patency [Fig. S5]. Along with our lipid data, we thus propose that patency is the mechanism by which the St 10a oocyte accumulates lipids under ecdysone regulation.

Endosymbionts also enter the oocyte from the maternal hemolymph in mid-vitellogenesis (*16)*, and we asked whether patency assists in this vertical transfer. Staining egg chambers for the endosymbiont *Spiroplasma pulsonii*, we found that some bacteria are present between the follicle cells prior to the onset of patency, while heavily populating the TC gaps at St 10a and 10b *(16)* [Fig. 6(A-B”)]. In FE lacking patency, *S. poulsonii* was still detected between the follicle cells as in earlier stages of vitellogenesis [Fig. 6(C-D”)]. These data suggest that patency is an advantage rather than a dependence for vertical transmission.

**Fig. 6.**
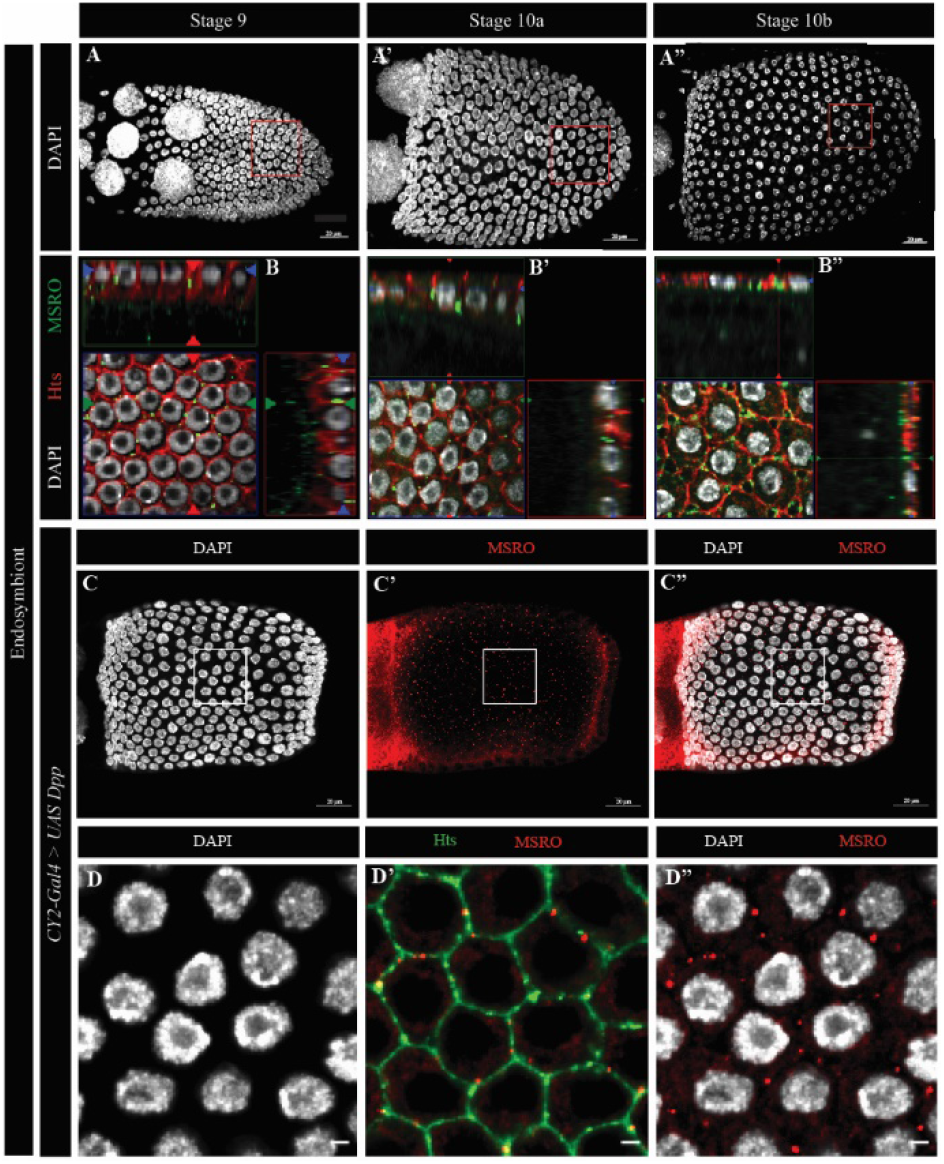
Vertical transfer of the endosymbiont *Spiroplasma pulsonii* across the FE between stages 9 and 11. (A-A”) Projections ofDAPI stained Stage 9 - 11 egg chambers. Red boxed are enlarged in (B-B”), and ortho views are provided for top and cross-sectional views. MSRO marks the bacteria, Hts marks the membrane, and DAPI stains the nuclei. (B) The bacteria are seen in the TCs of the FE at St 9 when they are still intact, as well as within the cells. (B’-B”) Bacteria are still present within the FE, and now occupy patent TCs at St 10a and 10b. (C-D”) MSRO staining in *UAS-Dpp* expressing FE shows the endosymbiont between the cells when the TCs remain intact, similar to WT St 9 with intact TCs.

### *D. simulans* FE presents similar spatiotemporal patterns of patency

To determine whether the temporal range and spatial pattern of patency are unique to *D. melanogaster*, we examined the ovaries of the closely related species *Drosophila simulans*. The FE of only St 10a and St 10b egg chambers have patent TCs in *D. simulans* as well [Fig. 7(A-D”’)]. The oocyte associated FE of St 10b egg chambers in *D. simulans* also presented a similar spatial pattern of patency as in *D. melanogaster,* with intact TCs in the dorsal anterior region [Fig. 7(E-F”)]. Together, our data indicates that the temporal and spatial patterns of patency could be evolutionarily conserved.

**Fig. 7.**
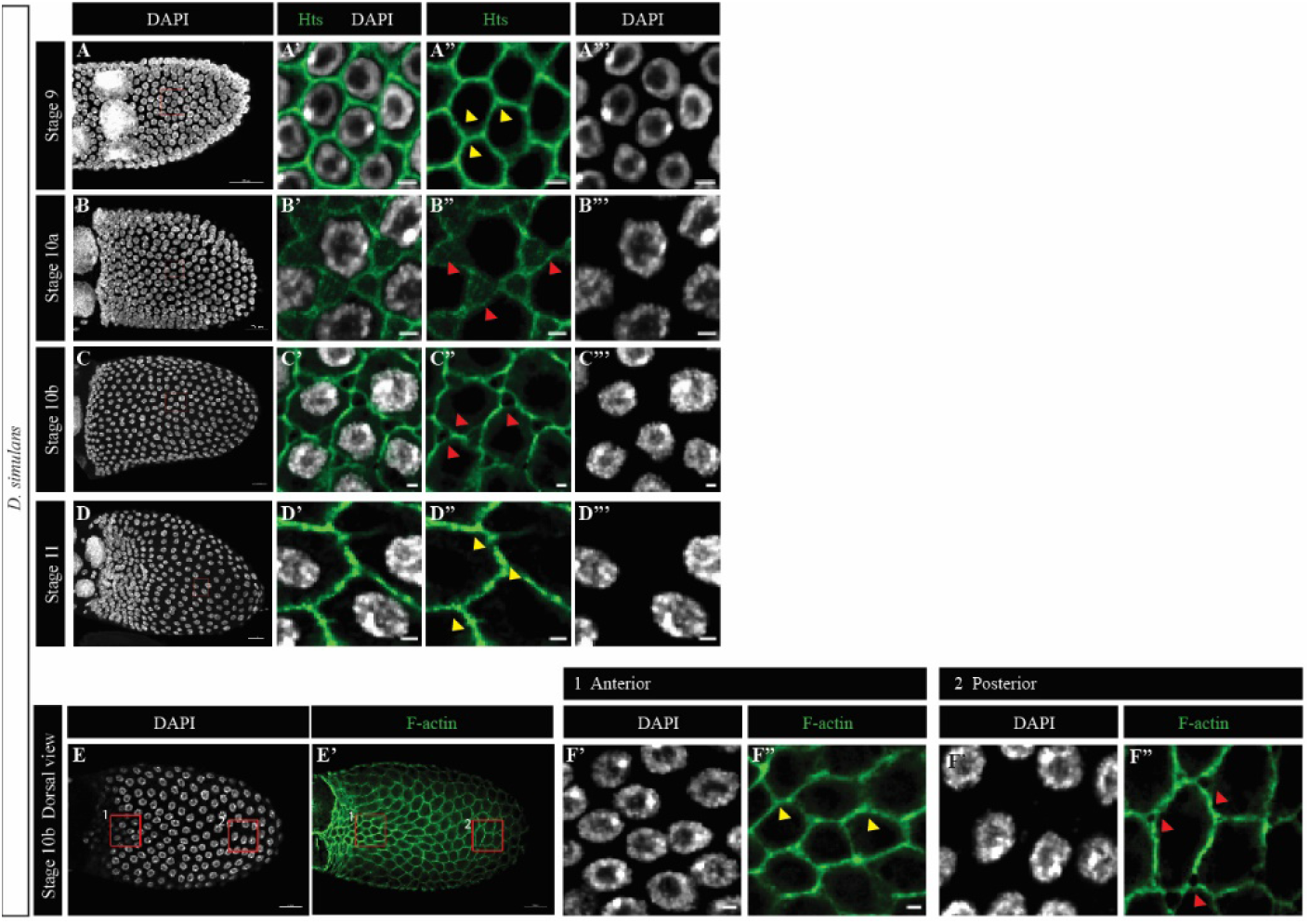
Temporal range and spatial pattern are conserved in *Drosophila simulans.* (A-D’”) Temporal range of patency in *D. simulans.* St 10a (B-B”’) and St 10b (C-C”’) show TC gaps characteristic of patency, while St 9 (A-A”’) and St l l(D-D’”) have intact TCs. (E-F”) Spatial pattern of patency in *D. simulans* is similar to that in *D. melanogaster.* (E-E”) Dorsal view ofa Stage 10b egg chamber in *D. simulans.* Box 1 in the dorsal anterior is expanded in (F-F’) showing intact TCs (yellow arrowheads). Box 2 in the dorsal posterior is expanded in (F”-F”’) showing TC gaps (red arrowheads).

**Fig. 8.**
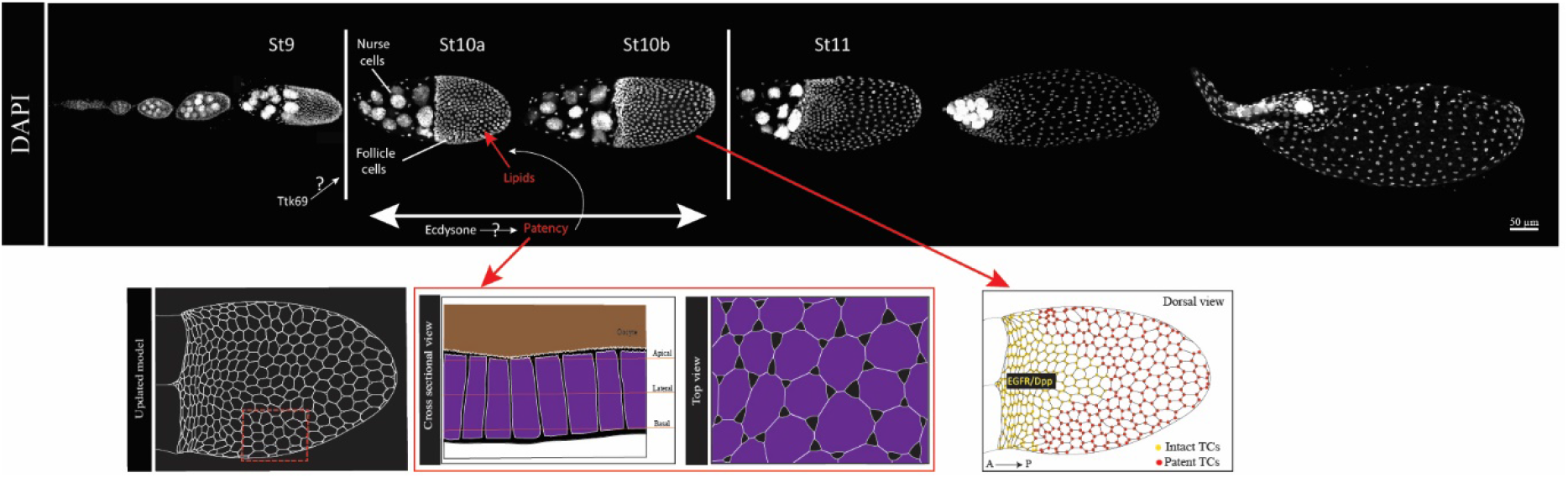
Model of follicular patency regulation in *Drosophila melanogaster.* DAPI stained egg chambers as arranged in an ovariole, depicting gennarium to St 13. Patency is observed in the Oocyte-associated follicular epithelium as gaps between the cells from the basal to the apical domain, illustrated in the cross-sectional and top-down views. The strict temporal range of patency (StlOa - 10b) is regulated by Ttk69, and the mechanistic details are yet to be discovered. We also found that Ecdysone signaling - which incidentally becomes activated in the FE at St 1Oa *(43)* - generates patency, and patency in tum is required for lipid uptake by the oocyte at St 10a. At stage ! Ob, the dorsalizing signals EGFR and Dpp create a spatial pattern of patency, illustrated as ‘intact TCs’ at the dorsal anterior, and ‘patent TCs’ in the rest of the FE.

## Discussion

In this study, we report that *Drosophila melanogaster* exhibits a phase in oogenesis when the oocyte-associated follicular epithelium develops ‘gaps’ that extend across the monolayer, creating a breach in the epithelium, a process referred to as ‘patency’. The FE, whose primary function is to maintain an intact shield around the egg chamber, is developmentally perforated with TC gaps essential for lipid transport. We illustrate spatiotemporal regulation of patency on a global scale by the ecdysone signal, at a local level by the axial patterning signals of the FE (Dpp and EGFR), and at a transcriptional level by Ttk69. These signals together regulate TC openings to allow trans-epithelial transport of lipids [model in Fig. 8].

The use of *CY2-Gal4* to drive the expression of genes vital for vitellogenic progression allowed us to tease out patent-stage specific effects of the regulators on the FE. For instance, our data showing the requirement of the Ecdysone signal for the formation of patent TCs [Fig. S5] suggests an important new role for the steroid hormone both temporally and functionally. In several insects such as *R. proxilus*, *L. migratoria*, and *T. castaneum*, the sesquiterpenoid juvenile hormone (JH) has been reported to be a major regulator of patency (*27–29*). A complex interaction between the two hormones coordinates vitellogenic progression in *Drosophila* (*30, 31*). It is, therefore, possible that patency in *Drosophila* could also be affected by JH, as well as its interaction with Ecdysone. Although Ttk69 is placed directly downstream of Ecdysone activation in later oogenesis *(22)*, our data suggests a more complex interaction for the regulation of patency.

While characterizing the junctions across patent stages, we observed the presence of Gli at the cessation of patency. Gli, although a central component of TCJ septate junctions *(19)*, is only one component of the TCJs in epithelia. Other known TCJ proteins and assemblers include Anakonda (Aka), the master regulator of Gli expression and localization *(32)*, Sidekick, the tricellular junction adherens junction protein (*33–35*), and M6, the TCJ protein required for oogenesis *(36)*. It is possible that these and other TCJ proteins might be involved in the creation, maintenance, and termination of patency in the FE. An assessment of the chronological order of appearance, localization, and expression levels over the course of patency, in addition to functional studies, will help delineate the mechanistic details of patency in the FE.

A previous report on tissue remodeling via a non-canonical secretion pathway aids in closing ‘open Zones of contact (ZOCs)’, and we propose that these open ZOCs are indeed the patent TCs we have characterized here *(37)*. Furthermore, our *D. simulans* data show that the temporal range of patency and its spatial pattern could be evolutionarily conserved. Further analysis of patency in other insects will ascertain whether the spatiotemporal patterns are evolutionarily conserved, specifically in meroistic ovaries similar to those of *D. melanogaster* and *D. simulans*. Altogether, this study creates a platform for further research into epithelial behavior, TCJ assembly and dynamics, and trans-epithelial transport mechanisms.

## Methods

### *Drosophila* strains and culture

All *Drosophila* stocks were maintained and crossed at 21–22°, unless otherwise indicated. The w[118] strain was used as the wild-type control. To control the temporal and regional gene expression targeting (TARGET) system, temperature-sensitive Gal80 (Gal80^TS^) under the ubiquitous tubulin driver was used to regulate the upstream activating sequence (UAS)-transgene expression by altering temperatures. *Tj-Gal4* and *CY2-Gal4* were used to drive UAS transgene expression in combination with Gal80^TS^. The crosses involving Gal80^TS^ were crossed and maintained at 18°C, and progeny were kept at 29°C for 48hrs prior to dissection.

Tj-Gal4 is used to drive transgenes under UAS control in all the follicle cells across oogenesis. Transgenes that disrupt vitellogenic progression were used with *CY2-Gal4*, that drives transgenes under UAS control in only the oocyte associated follicle cells starting at low levels at St 9, and at high levels from St 10a [Fig. S4.(C-E’)].

The following stocks were used: w[118] (Bloomington Drosophila stock Center (BDSC) #5905), *UAS Gli RNAi* (RNA interference) (BDSC #58115), *UAS ttk RNAi* (BDSC #27325), *UAS Dpp* (BDSC #1486), *Tj-Gal4* (BDSC #4775), *CY2-Gal4* (a gift from Dr. Berg), a*lphaTub84B-Gal4, UAS-mCD8:GFP* (BDSC#30030), *UAS EGFR DN* (BDSC #5364), *10XUAS-IVS-myr::td Tomato* (BDSC #32221), *UAS EcR B1 DN* (BDSC #6872), *D. Simulans* (a gift from Dr. Houle).

### Immunostaining and fluorescence microscopy

Ovaries were dissected, fixed, and stained with antibodies as described previously (*38, 39*). The following primary antibodies were used: Hu li tai shao (Hts) (1:5, 1B1), E-cadherin (Ecad) (1:20, DCAD2), Discs large (Dlg) (1:50, 4F3), and Fasciclin3 (Fas3) (1:15, 7G10) all from the Developmental Studies Hybridoma bank (DSHB), Laminin (gamma 1) (1:200, ab47651, Abcam), Gliotactin (1:50, a gift from Dr. Auld), MSRO (1:200, a gift from Dr. Lemaitre), Ttk69 (1:200, a gift from Dr. Xi). Alexa Fluor 488- or 546-conjugated goat anti-mouse and anti-rabbit secondary antibodies (1:400; Molecular Probes, Eugene, OR) were used. Phalloidin-546 (Invitrogen) was used to stain F-actin. Nuclei were labeled with DAPI (1:1000). Images were captured with Zeiss LSM 800 confocal microscope. Zeiss and ImageJ software were used for image analyses and processing.

Detergent-free protocol was modified from the standard protocol above, with all the same reagents prepared without Tween-20.

#### Lipid staining with Nile red and BODIPY

Ovaries were dissected in PBS and fixed in 4% PFA for 15 minutes. Following 2 washes with PBT, the ovaries were incubated in Nile red (TCI America, 7385-67-3) (0.002% dye diluted in PBT, adopted from 29) or BODIPY 493/503 (ThermoFisher Scientific, Cat No. D3922) (Stock: 1mg/ml BODIPY in absolute ethanol, Working: 1:500 in PBS) in dark for 30 minutes. Following 2 PBT washes, the ovaries were either stained with antibodies, or with DAPI and then mounted onto slides.

### Transmission Electron Microscopy

The samples for TEM were prepared as previously described (*40, 41*). Tissue samples were fixed with Karnovsky's Fixative. After three rinses with 0.1 M sodium cacodylate buffer, samples were embedded in 3% agarose and sliced into small blocks (1mm3), rinsed with the same buffer three times and post-fixed with 1% osmium tetroxide and 0.8 % Potassium Ferricyanide in 0.1 M sodium cacodylate buffer for one and a half hours at room temperature. Samples were rinsed with water and en bloc stained with 4% uranyl acetate in 50% ethanol for two hours. They were then dehydrated with increasing concentration of ethanol, transitioned into propylene oxide, infiltrated with Embed-812 resin and polymerized in a 60°C oven overnight. Blocks were sectioned with a diamond knife (Diatome) on a Leica Ultracut 7 ultramicrotome (Leica Microsystems) and collected onto copper grids, post stained with 2% aqueous Uranyl acetate and lead citrate. Images were acquired on a JEM-1400 Plus transmission electron microscope equipped with a LaB6 source operated at 120 kV using an AMT-BioSprint 16M CCD camera.

## Data Availability

All data generated or analyzed during this study are included in this manuscript and the supplementary information files.

## Acknowledgements and funding

We thank Drs. V. Auld, C. Berg, D. Houle, B. Lemaitre, R. Xi, Developmental Studies Hybridoma Bank (DSHB, USA), and Bloomington Drosophila Stock Center (BDSC, USA) for antibodies and fly stocks. We acknowledge the assistance of the UT Southwestern Electron Microscopy Core Facility for acquiring TEM images. We thank members of the Deng laboratory for valuable discussions and support, and P. Michael Albert II for assistance with editing the manuscript. W.-M. D. is supported by NIH R01 GM072562, R01 CA224381, R01 CA227789 and NSF IOS-1552333.

## Author contributions

Conceptualization, methodology, validation, investigation, visualization, project administration: S.R., W.-M.D.; Writing - original draft: S.R.; Writing - review & editing: S.R., W.-M.D.; Supervision, funding acquisition: W.-M.D.

## Materials and Correspondence

All data is available in the manuscript or the supplementary materials. Correspondence and material requests should be addressed to Wu-Min Deng at.

## Supplementary Information

Figures S1-S5

