## Supplementary Material for "Developmental regulation of oocyte lipid uptake via ‘patent’ follicular epithelium in *Drosophila melanogaster*"

**This file includes:**

Figs. S1 to S5
Captions for Data S1 to S5

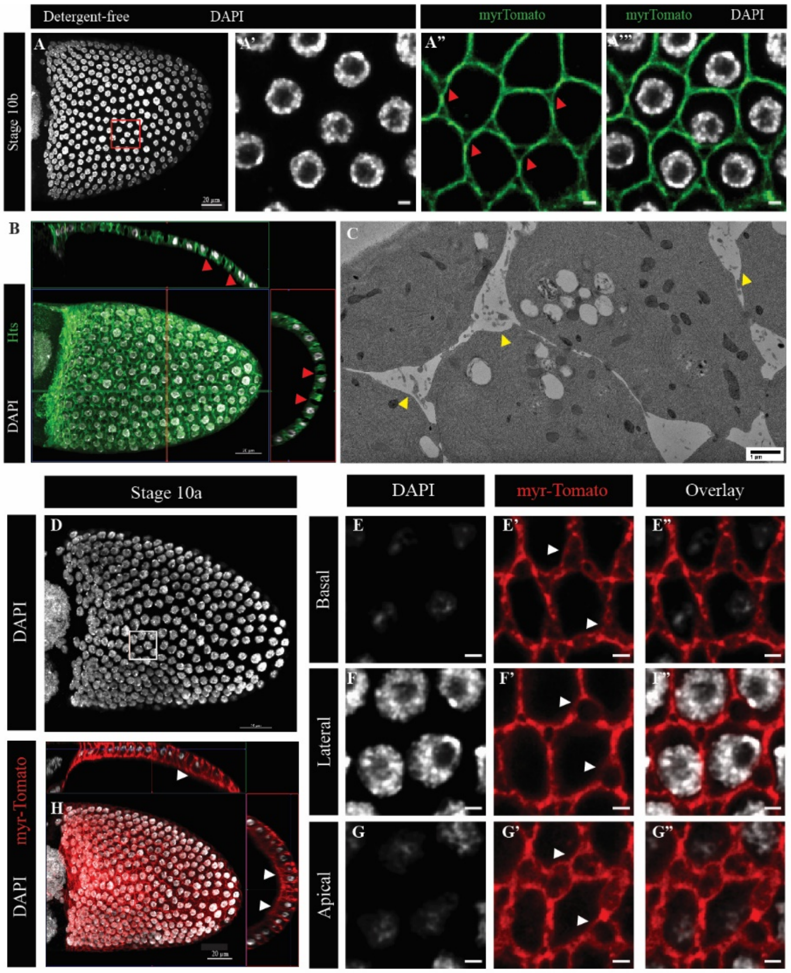


**Fig. S1.** Confirmation of TCJ gaps in *Drosophila* FE. (A) Stage 10b egg chamber expressing *Tj-Gal4*, *10XUAS-myr-Td-Tomato*, fixed and mounted using a detergent-free protocol (see materials and methods). Red box is enlarged in (A'-A''') showing patent TCJs even when detergent-free reagents are used to process the egg chambers. (B) Ortho view of a projection of a St10a egg chamber showing gaps extending across the FE monolayer (red arrowheads) Hts marks the membrane. (C) TEM image of a St10a egg chamber showing unidentified material in the gaps (yellow arrowheads). (D-G") Stage 10a oocyte-associated FE expressing *10XUAS-myr-Td-Tomato*. White box is expanded in (E-G"). (E-E", F-F", G-G") Basal, lateral, and apical view, respectively. (H) Ortho view of a projection of a St10a *10XUAS-myr-Td-Tomato* expressing egg chamber, showing gaps extending across the FE monolayer (white arrowheads). DAPI marks nuclei.

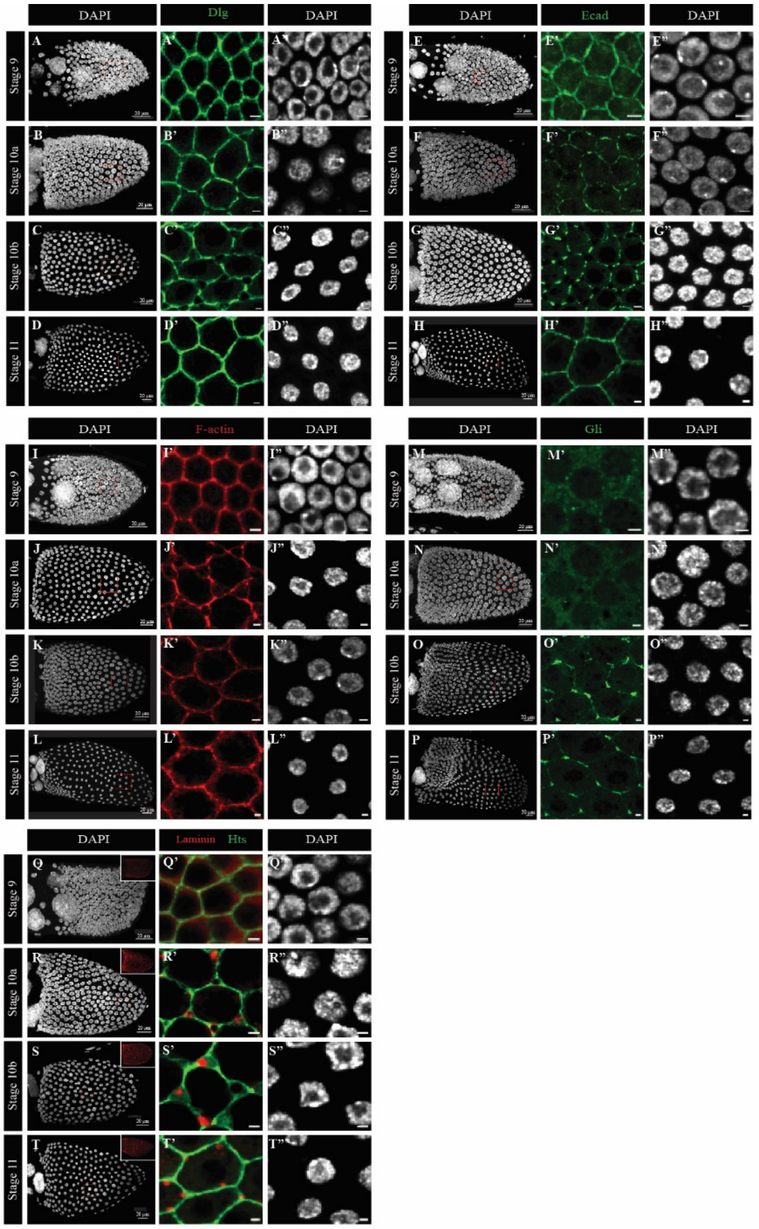


**Fig. S2.** Characterization of adhesion and junction proteins over patency between stage 9 and 11. (A-D") Septate junction protein Dlg distribution before (A-A"), during (B-C") and after (D-D") patency. (E-H") Adherens junction protein E-cadherin (R-cad) is intact before patency (E-E"), is removed from the TCs in patent stages (F-G"), and reappears when patency is terminated (H-H"). (I-L") Cortical F-actin (Phalloidin) remains intact in patent epithelia and continues to line the TCs when patent, making phalloidin a good stain to visualize TC gaps. (M-P") The tricellular septate junction protein, Gliotactin (Gli), is not present at intact TCs prior to (St9, M-M"), or at the onset of patency (St10a, N-N") but appears at the TCs at stage 10b (O-O") and remains after the termination of patency (P-P"). (Q-T") Localization of the basement membrane component Laminin. Absent from TCJs at St9 (Q-Q"), Laminin appears and localizes in the gaps in patent FE starting at St10a (R-R") and St10b (S-S"), and is still present near the TCs even at stage 11 after the gaps disappear (T-T"). Insets in (Q-T) Laminin in the full FE from stage 9 - 11.Consistent staging of egg chambers was based on the characterization by Jia et al, 2016 *(31)*.

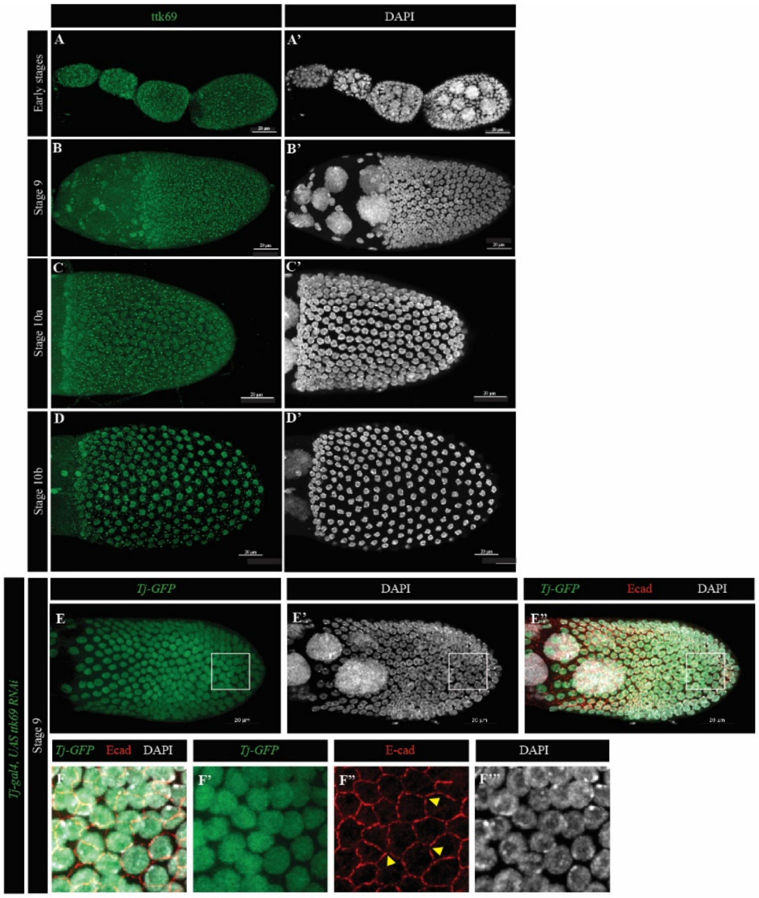


**Fig. S3.** Ttk69 pattern in WT FE. Fixed ovaries stained for Ttk69 with the antibody from the Xi lab (*42*). Ttk69 expression is observed in the nuclei of follicle cells even at early stages (A-A’), as well as at St9 (B-B’), St10a (C-C’), and St10b (D-D’). Punctate signals in the FE can be attributed to background noise. DAPI marks nuclei. (E-F””) Stage 9 egg chamber expressing *UAS-ttk69-RNAi* under *Tj-Gal4*. (E-E”) Projection of a St9 ttk69 KD egg chamber with E-cad immunostaining. White box is expanded in (F-F””). E-cad is lost from the TCs (yellow arrowheads). DAPI marks the nucleus in grey.

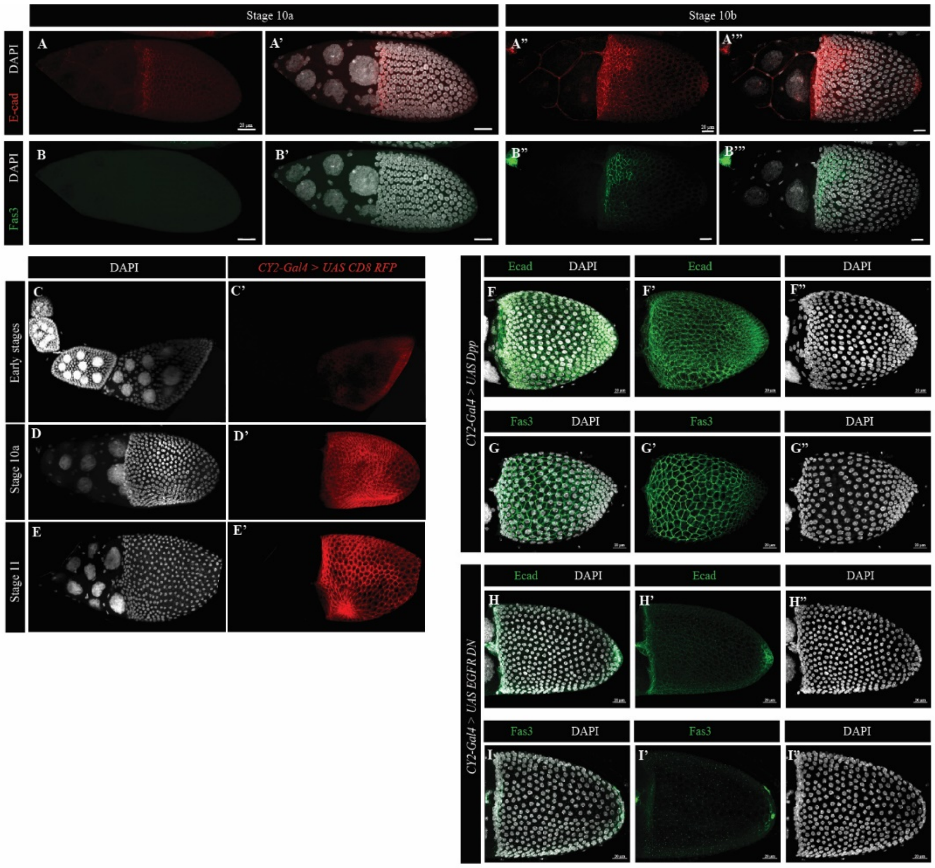


**Fig. S4.** Dorsal pattern of adhesion proteins E-cad and Fas3. (A-B’’’) WT egg chambers develop the distinct distribution of E-cad and Fas3 at the dorsal anterior from St10a (A-A’, B-B’ respectively) to St10b (A”-A’”, B”-B”’, respectively). (C-E”) *CY2-Gal4* expression pattern in *Drosophila* ovaries marked by *UAS-CD8:RFP* on the membrane. Expression in the oocyte-associated FE begins at St9 (C-C”) and continues through 10a (D-D”) and St11 (E-E”). (F-G”) Egg chambers expressing *UAS-Dpp* under *CY2-Gal4* driver. Expansion of the domains with high E-cad (F-F”) and the Fas3 (G-G”) at stage 10b. (H-I”) Egg chambers expressing a dominant negative form of EGFR under UAS control using the *CY2-Gal4* driver. The dorsal pattern of E-cad (H-H”) and Fas3 (I-I”) are lost in the FE at stage 10b.

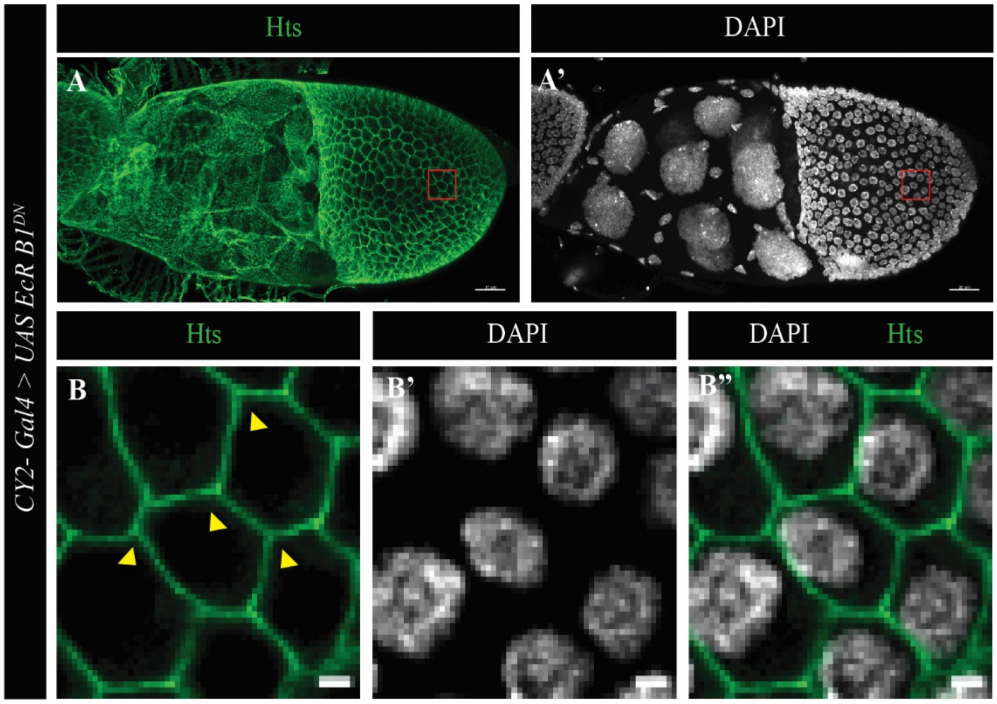


**Fig. S5.** Ecdysone signaling is required for patency. (A-A') St10a egg chamber expressing a dominant negative form of the ecdysone receptor EcR B1 under *CY2-Gal4*. Red box is enlarged in (B-B"), showing intact TCs (yellow arrowheads). Hts marks the membrane, DAPI marks nuclei.
